# Effectiveness of treatments for firework fears in dogs

**DOI:** 10.1101/663294

**Authors:** Stefanie Riemer

## Abstract

An online questionnaire survey investigated (1) management and (2) treatment methods for firework fears in dogs employed by dog owners and their (perceived) effectiveness. A PCA on data from 1225 respondents revealed four management strategies (i.e. interventions during firework exposure): the principal components *“Environmental modification”* (e.g. providing a hiding place, keeping windows and blinds closed, and playing music), *“Feed/Play”* (providing the dog with chews, play and food during fireworks in general, as well as contingent on loud bangs), *“Alternative”* (use of calming nutraceuticals, pheromones, herbal products, homeopathic products, Bach flowers, and essential oils) and *“Interaction”* (allowing body contact, petting and talking to the dog when loud bangs occurred). To explore possible effects of these management methods on fear development, the components were correlated with a score for fear progression. Of the four components, only *“Feed/Play”* was statistically associated with an improvement in fear responses to fireworks. To evaluate the effectiveness of various treatment strategies, owners were asked to select from a range of options which interventions they had used and whether they considered them as effective. With prescription medication (N=202), improvements were noted by 69% of owners, with high success rates reported for the most frequently prescribed drugs, alprazolam (91%) and Sileo® (74%). While individual products were not evaluated, the reported success rates for the categories “pheromones” (N=316), “herbal products” (N=282), “nutraceuticals” (N=211), “essential oils” (N=183), “homeopathic remedies” (N=250) and “Bach flowers” (N=281) were all in the range of 27-35%, which is not higher than would be expected based on a placebo effect. Pressure vests were deemed as effective by 44% of respondents (N=300). Counterconditioning (providing desirable stimuli after the occurrence of noises) was the most successful training technique according to the owners (N=694), with a reported effectiveness of over 70%. Relaxation training (N=433) was reported to be almost as successful at 69%, while noise CDs (N=377) were effective in 55% of cases. Thus, counterconditioning, relaxation training and anxiolytic medication appear to be the most effective strategies in the treatment of firework fears in dogs. On this basis it is recommended that ad-hoc counterconditioning and relaxation training should complement the standard behavioral technique of desensitization/ counterconditioning with noise recordings.

**Highlights:**

- An online survey on treatment options for firework fears in dogs was performed
- Feeding or playing with dogs during fireworks was associated with fear improvement
- Success was highest for ad-hoc counterconditioning, relaxation training and medication
- Success was similar for pheromones, nutraceuticals and other alternative products
- Success rates for these alternative products are consistent with a placebo effect

## Introduction

Fear responses to fireworks and other loud noises (often referred to as ‘noise phobia’, ‘noise sensitivity’, ‘noise reactivity’, ‘noise aversion’ or ‘noise stress’) represent a significant welfare problem for pet dogs. According to surveys, up to half the pet dog population react fearfully to fireworks (Blackwell et al., 2013; Riemer, 2019; Storengen and Lingaas, 2015; Tiira et al., 2016), and one study indicated that over 15% of fearful dogs require several days or longer to recover behaviorally from a firework event, with over 3% exhibiting behavioral changes for weeks or months (Riemer, 2019). Thus, identifying effective interventions to prevent and treat firework fears in dogs is of wide concern. Owners’ awareness of the issue seems to be increasing (c.f. Riemer, 2019), and a wide range of products are marketed as alleviating firework fears in dogs, ranging from pheromones to special garments and from homeopathy to prescription medication. However, as has been pointed out previously, many products lack scientific evidence for their effectiveness, and even most published studies in the field are based on low sample sizes and often lack placebo treatments (Sherman and Mills, 2008). Subsequently I give a brief overview over approaches to treating noise fears in dogs.

### 1) Behavioral techniques

#### Management

In order to minimize adverse experiences for the animal, management measures are important when exposure to fearful stimuli cannot be avoided (Sherman and Mills, 2008). Commonly given management advice for noise fears includes providing a safe place to retreat, ideally by associating it with positive experiences prior to any firework events, darkening the room putting music or white noise on, ignoring fearful behavior, distracting the dog with games, training or food, and refraining from any punishment (Mills, 2005; Pike et al., 2015; Sherman and Mills, 2008). However, management is likely not to be sufficient to generate long-lasting improvement, and behavior modification techniques, often with adjunctive use of pheromones or medication, are usually recommended (Sherman and Mills, 2008). Given that fears and phobias can significantly compromise welfare and become severe, wherever possible clinicians should aim to treat fears and attempt to resolve the problem, rather than relying on management only (Horwitz and Mills, 2012).

#### Systematic desensitization and counterconditioning

‘Desensitization” can be defined as “gradual and controlled exposure to the stimulus so as to extinguish the manifestations of fearful behavior” (Horwitz and Mills, 2012, p. 177). It involves repeated exposure to the feared stimulus, but at such a low intensity that it elicits no distress response. For instance, a noise recording can be played at a very low level. While the dog might initially alert to it, it should learn to ignore it with repeated exposure. Gradually, over the course of multiple sessions, the volume can be increased, as long as the dog does not show any distress upon exposure (McMillan, 2019). Generally, this approach is most effective in alleviating fear responses when combined with counterconditioning (McMillan, 2019).

‘Counterconditioning’ usually refers to pairing desirable stimuli such as food or play with the fear-eliciting stimulus (‘respondent’ or ‘classical counterconditioning’) in order to reduce the animal’s distress through replacing the association with a positive one (McMillan, 2019). Delivery of the incentive is contiguous only with the feared stimulus, regardless of the animal’s behavior. A behavioral change should follow as a result of a change of the animal’s emotional or motivational state (McMillan, 2019).

Some authors also describe the process of ‘operant’ or ‘instrumental counterconditioning”, defined as “reinforcing a substitute behavior that is incompatible with the unwanted behavior” (Horwitz and Mills, 2012, p.217). In contrast to ‘classical counterconditioning’, upon exposure to the aversive stimulus, the animal is required to perform an operant response in order to receive an incentive (McMillan, 2019). Based on early research, classical counterconditioning is considered as superior to operant counterconditioning when aiming to change negative emotional states (McMillan, 2019). Hereafter, I use the classic definition of the term ‘counterconditioning’ as meaning that the feared stimulus is paired with an appetitive/positive outcome (e.g. food) with the goal of decreasing the fear response over repeated pairings and replacing it with an appetitive response (c.f. Newall et al., 2017).

Counterconditioning is most effective when combined with desensitization, i.e. the feared stimulus is initially presented at a very low level and followed by an incentive. Subsequently, this process can be repeated, whereby stimulus intensity can be gradually increased. Nonetheless, using a desensitization approach in combination with counterconditioning is not always possible as not all fear-inducing stimuli can be under complete human control, and some fear improvements can still be achieved using counterconditioning when stimuli are presented at full intensity. Crucial to the success of counterconditioning is identifying a highly valued incentive (McMillan, 2019).

The use of noise recordings for systematic desensitization and counterconditioning (abbreviated as DSCC) is the most commonly advised behavioral technique in the treatment of noise fears (e.g. (Horwitz and Mills, 2012; Levine et al., 2007; Levine and Mills, 2008; Mills et al., 2003; Sherman and Mills, 2008). Studies indicate that most owners perceive an improvement in their dogs’ firework fears after following a DSCC program using noise CDs and are very satisfied with the treatment (Levine et al., 2007; Levine and Mills, 2008; Sheppard and Mills, 2003), even up to one year later (Levine and Mills, 2008). However, one study reports that no effect of treatment could be discerned using objective behavioral measures when dogs were exposed to a novel CD recording in the clinic setting (Levine et al., 2007), and this finding was similar to that of a previous study on the use of noise CDs (in combination with medication) on thunderstorm fear in dogs: Despite owner-reported improvement, no change in behavioral signs was noted during post-treatment exposure to a thunderstorm recording in the clinic (Crowell-Davis et al., 2003).

#### Relaxation training

Training animals to relax on cue is a less commonly recommended tool in behavior modification (but see Mills, 2003; Overall, 2013; Talegón and Delgado, 2011). Relaxation may be achieved by different approaches. Horwitz and Mills (2012) suggest to firstly, induce relaxation by massage or long strokes. Secondly, this can be associated with a word in order to classically condition a calm physiological state. After successful conditioning, this cue can then be used to induce relaxation during stressful events even without the need to massage continuously (Horwitz and Mills, 2012). While this method relies on classical conditioning, an operant ‘protocol for training relaxation’ is given in Overall (2013). Here, the dog is progressively rewarded for behaviors, facial or bodily expressions consistent with relaxation, and for remaining still for increasing amounts of time and in the face of gradually increased distractions (Overall, 2013b). To what extent such training is effective in reducing fear or anxiety in dogs has not been investigated to date.

### 2) Pharmacological interventions

#### Prescription medication

While benzodiazepines (e.g. Alprazolam), MAOIs, SSRIs, trazodone and the α2-adrenoceptor agonist clonidine can be indicated in the medical management of noise fears in dogs (Horwitz and Mills, 2012; Overall, 2013), only a handful of studies have investigated the effects of medications on noise fears in dogs, and surprisingly few were placebo-controlled. Two recent high-quality randomized, double-blind, placebo-controlled, clinical field studies demonstrated a high effectiveness of Sileo® Dexmedetomidine oromucosal gel (Korpivaara et al., 2017) and imepitoin (Engel et al., 2019), respectively. For Sileo®, 72% of owners reported a good excellent treatment effect compared to only 37% in the placebo group (Odds ratio in favor of Sileo®: 3.4; 95% confidence interval 1.95 to 5.99; p<0.0001). This medication is already licensed for noise fears in dogs in the European Union at the time of publication (European Medicines Agency, 2015). For imepitoin, around two thirds of owners considered the treatment effect good or excellent treatment compared to approximately a quarter of owners from the placebo group (Odds ratio in favor of imepitoin 4.7; 95% confidence interval 2.79-7.89; p<0.0001), which was also corroborated by highly significant differences between the two groups in an additional measure, (owner-reported) anxiety scores (Engel et al., 2019).

Only a few small, non-placebo controlled studies exist for the effect of other types of medication on the expression of noise fears in dogs. Two small open-label studies indicated that trazodone and clonidine, respectively, were effective in alleviating storm or noise phobia in dogs where other treatments (including other medications) had failed (Gruen and Sherman, 2008; Ogata and Dodman, 2011). A small prospective open clinical trial investigated the effect of a combination of daily clomipramine, added alprazolam before storms, and desensitization/ counterconditioning at home with a thunderstorm audio recording on the behavior of dogs affected by fear of thunderstorms. While almost all caregivers indicated an improvement in their dogs, this could not be confirmed when exposing dogs to the audio recording in the clinic, although it is not clear whether post-treatment recordings were made under the influence of alprazolam (Crowell-Davis et al., 2003). Another small retrospective study indicated that diazepam was considered very or somewhat effective by 67% of owners, but many owners discontinued the treatment due to the occurrence of side effects (Herron et al., 2008).

### 3) Alternative products

#### Pheromonatherapy

Pheromonatherapy utilizes analogues of animal pheromones, chemical signals normally involved in intraspecific communication that are processed by the vomeronasal organ and are assumed to have an intrinsic effect on the emotional processing of animals (Mills et al., 2012). Dog appeasing pheromone (DAP, Adaptil®) mimics a pheromone produced by lactating bitches after parturition that is believed to instill a sense of well-being in the puppies. It is used in veterinary behavioral medicine as a calming agent also for adult dogs (Mills et al., 2012).

An open-label follow-up study indicated that the use of Dog Appeasing Pheromone plug-in diffusers resulted in a high owner satisfaction level and reported improvement in dogs’ clinical signs during firework exposure (Sheppard and Mills, 2003), although the inclusion of behavioral advice in the treatment plan does not allow to distinguish effects of behavioral modification and pheromones (c.f. Frank et al., 2010). DAP was also used as in the above-cited studies in combination with noise CDs (Levine et al., 2007; Levine and Mills, 2008). One study suggested that the use of DAP and noise CDs in combination had higher owner-reported success rates than when either intervention was used alone (Mills et al., 2003). While the above studies were unblinded and did not include a placebo group, a blinded parallel-group placebo controlled study on the effects of DAP collars on the behavior of laboratory beagles during playbacks of thunderstorm recordings concluded that DAP is of potential benefit as an adjunct to a behavior management program (Landsberg et al., 2015a). In this study, observations of active (such as startling, scanning, bolting, pacing, running, circling, digging, climbing, jumping and barking) and global fear scores were significantly lower in the pheromone-treated group compared to the placebo group. No significant group differences emerged in passive scores (such decreased activity, freezing, lower body postures, panting, trembling, salivating, lip licking etc.). Hiding in a hiding box occurred significantly more in placebo-treated subjects (Landsberg et al., 2015a).

#### Nutraceuticals

Given that nutraceuticals do not underlie the same stringent approval processes as medications and therefore enter the market much more easily, many non-prescription nutraceuticals are available for treating fear and anxiety in dogs (Orlando, 2018). However, little published research on their effects exists, especially in regard to noise fears. For example, Zylkene (alpha-casozepine) may be recommended to aid with noise fears although it has not specifically been tested for this indication (Horwitz and Mills, 2012). The effect of L-theanine on fear of thunder in dogs was investigated in a small open-label trial. However, a high drop-out rate (eight of originally 26 subjects could not be included in the analysis due to early withdrawal or failure to complete the paperwork) hinders drawing valid conclusions. For the remaining 18 dogs, a highly significant reduction in owner-reported global anxiety scores and time to return to baseline following a storm was found, and treatment success (defined as “an improvement in the behavior score after the 5th storm compared with baseline of at least 1 in 50% of the behaviors identified, with no behavior getting worse” was achieved in 12 of the 18 dogs (Pike et al., 2015). A fish protein supplement appears to have beneficial effects on cortisol reactivity in dogs exposed to a thunderstorm recording (Landsberg et al., 2015b).

Many owners tend to prefer to try “natural treatments” – such as herbal remedies, homeopathy or essential oils – in the management of behavioral problems first, with medication being regarded as only a last resort (Notari and Gallicchio, 2008). However, for most of these products, there is either no peer-reviewed research at all, results are inconclusive or negative.

#### Herbal formulations

A placebo-controlled crossover study investigated the effect of Harmonease Chewable Tablets, containing a blend of extracts of *Magnolia officinalis* and *Phellodendron amurense*, on inactivity duration during thunder storm recordings in 20 beagle dogs (with higher inactivity interpreted as reflecting greater fear intensity). While an analysis of variance indicated no significant treatment or order effect, the number of dogs reducing levels of inactivity during thunderstorm recordings was higher in the treatment group (60%) than in the placebo group (25%) (DePorter et al., 2012). Note, however, that later studies found that many individuals react with an increase, rather than a decrease, in activity to thunderstorm recordings (Gruen et al., 2015; Landsberg et al., 2015a, 2015b).

#### Homeopathy

A double-blind placebo-controlled study on the use of a homeopathic remedy in the treatment of firework fears in dogs showed high owner-reported improvement rates in both the placebo group (65%) and the verum group (71%), with no significant differences in any measures between the two groups. While the owners were also given simply management advice on how to react during firework exposure, this is unlikely to explain the perceived high rates of success, with substantial placebo effects being more likely (Cracknell and Mills, 2008). As such, open-label studies should be interpreted with caution.

#### Aromatherapy (essential oils)

Odors, in particular essential oils, may be used in a therapeutic context to alleviate stress in animals (Mills et al., 2012). As pointed out in the BSAVA Manual of Canine and Feline Behavioral Medicine, there are suggestions that lavender and chamomile may promote calmness in shelter dogs, but no evidence is available in relation to noise fears (Horwitz and Mills, 2012).

#### Bach flowers

Although there is no science-based evidence for their effectiveness, Bach flowers may be advised by some animal behavior professionals based on owners’ preferences (Notari and Gallicchio, 2008). Moreover, like Dog Appeasing Pheromone, herbal formulations, homeopathic products and essential oils, Bach flowers are freely available over the counter and so do not require a consultation with a professional.

### 4) Pressure vests

Studies on humans and on cattle indicate that deep pressure and weighted vests, or pressure in a squeeze chute, respectively, have calming effects. While the mechanism of action is not clear, it is suggested that peripheral oxytocin in response to the skin contact may contribute to relieving stress, reducing blood pressure and heart rate (reviewed in Pekkin et al., 2016). So-called pressure vests, tight-fitting vests for dogs exerting deep pressure when worn, utilize this principle and are postulated to exhibit calming effects on pet dogs. While the available evidence is inconclusive, there are some indications of effects on some behavioral parameters and heart rate during thunderstorm or firework recordings that could be interpreted as reflecting lowered anxiety when wearing a vest (Fish et al., 2017; Pekkin et al., 2016). Also owners considered the vest to be of benefit during real-life thunderstorm events (Cottam et al., 2013). A review concluded that “pressure vests may have small but beneficial effects on canine anxiety and that habituating the dog to the vest, assessing for comfort and using repeatedly may improve the likelihood of any benefit” (Buckley, 2018). Several vests are commercially available, such as the Thundershirt, Anxiety Wrap and Lymed Animal™ Supporting Garments.

To conclude, there is a lack of high-quality evidence for the majority of commonly recommended interventions in the treatment of noise fears in dogs. Since almost all existing studies suffer from very low sample sizes, it was decided to take a cross-sectional approach here using an online internet survey targeting a variety of different interventions. Like most previous studies, the study is based on the owners’ assessment of their dogs’ wellbeing during noise exposure and the effectiveness of behavioral techniques or products used.

## Methods

An online questionnaire survey (in an English and a German version) was distributed to dog owners via our research group’s website and social media. The questions covered the owners’ consent for the use of their data, demographic data about the dogs (date of birth, sex, neuter status, breed, country, source of dog, age at acquisition), dogs’ health problems and other potential behavioral problems (Riemer, 2019). Two scores were used to rate the severity and progression of firework fears, respectively (see below). Furthermore, owners were asked to indicate which management and treatment options they had used by selecting a number of options and, for the latter, whether they considered these as effective.

### Welfare impaired score

As a measure of the severity of firework fears, participants were asked “Please rate your level of agreement with the following statement: The overall welfare of my dog is strongly compromised by fireworks”. This was rated on a 5-point Likert scale ranging from “disagree strongly”, “tend to disagree”, “partly/partly”, “tend to agree” to “agree strongly”. The ensuing “Welfare Impaired” score was converted into a numerical score from 1 (“disagree strongly”) to 5 (“agree strongly”) for the analysis.

### Fear progression score

The “Fear progression” score was based on the question “How has your dog’s fear of fireworks progressed in the last years?”, with the following response options “My dog was never afraid of fireworks”, “The fear has improved greatly”, “The fear tends to have improved”, “The fear has remained the same”, “The fear tends to have become worse”, “The fear has become much worse” or “I don’t know”. For analysis, this was converted into a score ranging from 1 (“The fear has become much worse”) to 5 (“The fear has improved greatly”), with “I don’t know” treated as missing data.

### Management strategies

Owners were asked to select which management approaches (i.e. strategies during firework exposure, independent of any preparatory interventions) they had used to help their dogs cope with fireworks from a list of options (Table 1). They were also asked to select how they responded to their dogs when hearing loud bangs (Table 1).

**Table 1.** Questions related to owners’ management of firework fears (i.e. strategies during firework events).

|  |  |
| --- | --- |
| Which measures do you take on<br>New Year's eve/ on days with<br>fireworks? (Please select all<br>applicable answers). |  |
|  | Nothing |
|  | I create a safe hiding place for my dog |
|  | I close the curtains/ window shutters |
|  | I play loud music to drown out the noise |
|  | I put a thundershirt/ anxiety wrap on my dog |
|  | I offer chews to my dog |
|  | I play with my dog |
|  | I give food to my dog |
| How do you react yourself (to your<br>dog) when hearing loud bangs?<br>(Please select all applicable<br>answers). |  |
|  | I ignore him or her/ I don't pay attention to him or her |
|  | I pet him or her |
|  | I give him or her a treat |
|  | I play with him or her |
|  | I allow him or her to maintain body contact |
|  | I speak to him or her in a calming way |
| Have you administered medication or other calming products to your dog during the last firework event? | Yes - No |
| If yes, which ones? (Please select all applicable products). |  |
|  | Nutraceuticals (e.g. Zylkene, tryptophan etc.) |
|  | Pheromone products (Adaptil diffuser, spray or collar) |
|  | Herbal products |
|  | Homeopathic products |
|  | Bach flowers |
|  | Essential oils |
|  | Drugs (prescription medication available only from a veterinarian) |
|  | Other |

### Effectiveness of interventions

Owners were asked to select from a number of options which interventions to prevent or treat their dog’s fear of fireworks they had tried and whether they considered these as effective (response options: effective – not effective – not used; Table 2).

**Table 2.**
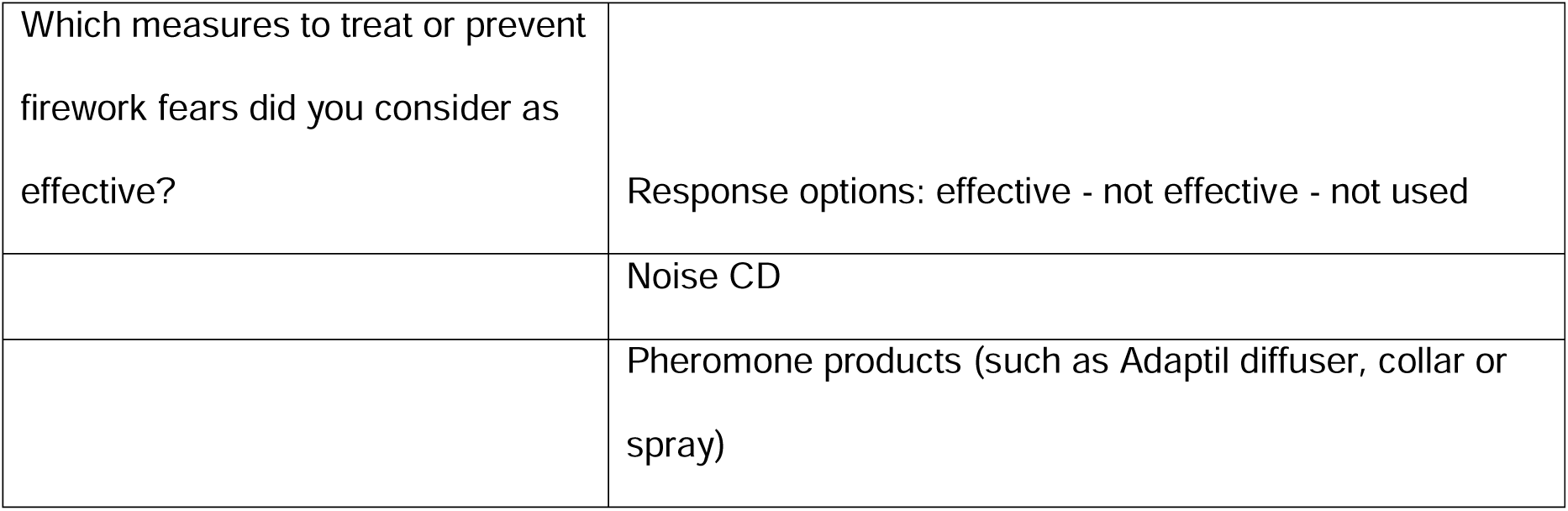
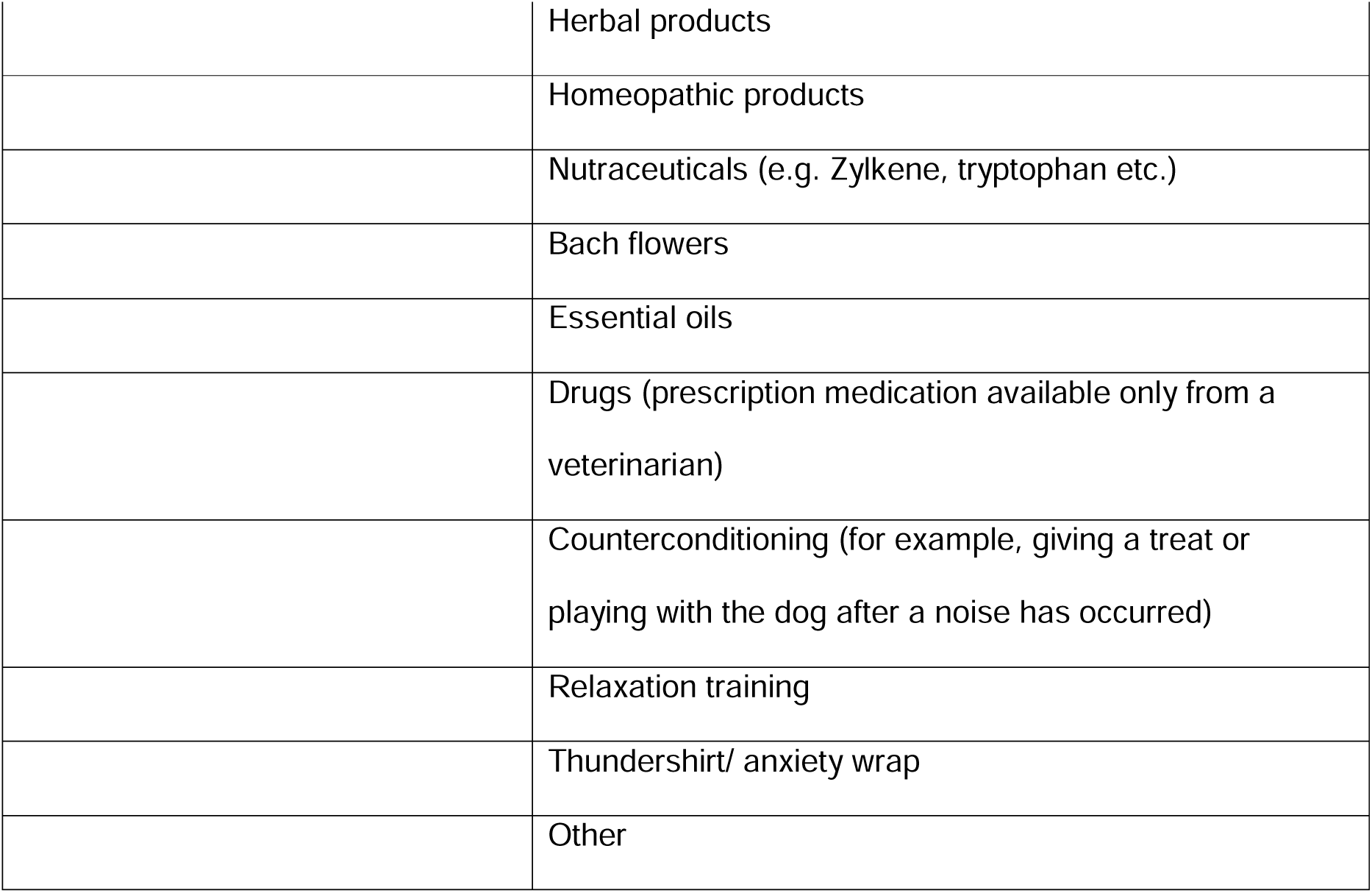
Question on the perceived effectiveness of interventions for management or training.

### Analysis

In order to identify patterns in management strategies, a non-linear principal components analysis (Linting et al., 2007; Linting and van der Kooij, 2012) or Categorical Principal Components Analysis (CATPCA) in IBM SPSS Statistics Version 23 (© IBM Corporation and its licensors 1989, 2015) was performed over the questions on owners’ management strategies during fireworks, including measures taken during New Year’s Eve/ on days with fireworks (e.g. creating a hiding place, closing windows, playing music etc.), how owners reacted to loud bangs themselves (e.g. ignoring the dog, petting the dog, giving the dog a treat etc.), and which types of products they had administered (e.g. pheromone products, prescription medication, pressure vests etc.). To assess relationships of management factors and fear development, the ensuing components were correlated with the Fear Progression Score using Spearman rank correlation tests, as requirements for parametric analysis were not met, using Statistica 6.1 (Statsoft Inc. 1984–2004).

For the specific questions on different products and training techniques used, the percentage of respondents considering a given intervention as effective was calculated from the total number of respondents that indicated having used this intervention.

## Results

A total of 1225 responses were analyzed, including 527 responses in English and 699 in German. Dogs were of various breeds or crosses and included 588 females (of which 430 neutered, 157 intact and one of unknown neuter status) and 637 males (of which 424 neutered, 207 intact and 6 chemically castrated). Over half the dogs in the survey (N=639) were given a rating of 3 or higher on the Welfare Impaired score and so are considered as fearful of fireworks (see (Riemer, 2019) for more details).

### Management strategies

Regarding owners’ management during firework exposure, the CATPCA yielded four factors (based on Eigenvalues >1 and Cronbach’s Alpha >0.7) that had no cross-loadings and accounted for 49% of the variance (Table 3).

**Table 3.** Results of a CATPCA on management strategies employed by owners during fireworks, with Varimax rotation. Loadings >0.4 are bolded.

|  |  | Environmental<br>Modification | Feed/ Play | Alternative | Interaction | Total |
| --- | --- | --- | --- | --- | --- | --- |
| Measures during firework events | No measures | **-0.718** | -0.210 | -0.087 | -0.256 |  |
|  | Hiding place | **0.654** | 0.049 | 0.148 | 0.217 |  |
|  | Windows closed | **0.803** | 0.104 | 0.100 | 0.100 |  |
|  | Music | **0.722** | 0.131 | 0.164 | 0.030 |  |
|  | Thundershirt | 0.351 | 0.077 | 0.400 | 0.044 |  |
|  | Chews | 0.238 | **0.684** | 0.038 | 0.003 |  |
|  | Play | 0.130 | **0.751** | -0.064 | 0.081 |  |
|  | Food | 0.074 | **0.749** | 0.071 | 0.086 |  |
| Reaction to bangs | Ignore | -0.057 | -0.071 | -0.021 | **-0.629** |  |
|  | Pet | 0.040 | 0.208 | 0.104 | **0.760** |  |
|  | Feed | -0.001 | **0.658** | 0.035 | 0.323 |  |
|  | Play | -0.035 | **0.718** | -0.005 | 0.193 |  |
|  | Body contact | 0.313 | 0.115 | 0.092 | **0.585** |  |
|  | Talk | 0.257 | 0.189 | 0.083 | **0.701** |  |
| Products used | Nutraceuticals | 0.244 | 0.024 | **0.443** | 0.030 |  |
|  | Pheromones | 0.248 | 0.005 | **0.586** | 0.021 |  |
|  | Herbal products | 0.123 | 0.059 | **0.729** | 0.000 |  |
|  | Homeopathy | 0.078 | -0.036 | **0.748** | 0.051 |  |
|  | Bach flowers | 0.112 | -0.057 | **0.645** | 0.094 |  |
|  | Essential oils | -0.044 | 0.053 | **0.580** | 0.086 |  |
|  | Prescription medication | 0.394 | -0.048 | 0.213 | 0.066 |  |
|  | **Cronbach's alpha** | **0.769** | **0.731** | **0.726** | **0.708** | **0.948** |
|  | **Eigenvalue** | **2.795** | **2.735** | **2.693** | **2.112** | **10.335** |
|  | **% of variance** | **13.309** | **13.021** | **12.825** | **10.058** | **49.214** |

The first component, termed ***“Environmental modification”*** had high positive loadings for providing a hiding place, keeping windows and blinds closed, and playing music and a high negative loading for not taking any measures. The use of a thundershirt or anxiety wrap and prescription medication also loaded > 0.35 on this component. The second component, ***“Feed/Play”***, had high positive loadings for providing the dog with chews, play and food during fireworks in general, as well as for feeding and playing with the dog in response to loud bangs. The third component, labelled ***“Alternative”*** had high positive loadings for the use of calming nutraceuticals, pheromones, herbal products, homeopathic products, Bach flowers, and essential oils. The fourth component, ***“Interaction”***, had high positive loadings for allowing body contact, petting and talking to the dog when loud bangs occurred, and a high negative loading for ignoring the dog.

Within those dogs with Impaired Welfare scores of 3 or higher, the principal component “Environmental Modification” was significantly positively correlated with Fear Progression (i.e. an increase in severity of firework fears, Spearman rank correlation test, Rho=0.181, N=566, p=0.00001) whereas the component “Feed/ Play” was significantly negatively correlated with Fear Progression (Spearman Rho=−0.16, N=566, p=0.00008). There was no relationship between Fear Progression and the components “Alternative” (Spearman Rho=0.026, N=566, p=0.527) and “Interaction” (Spearman Rho=−0.03, N=566, p=0.357).

### Owner-reported effectiveness of training techniques and products used

Noise CDs were considered as effective in alleviating dogs’ firework fears by 54.4% of owners who had used them (N=377; Figure 1). The reported effectiveness was 70.8% for counterconditioning (N=694) and 69.3% for relaxation training (N=433). A success rate of 44% was indicated for the use of a thundershirt/ anxiety wrap (N=300). For prescription medication, the reported rate of effectiveness was 68.9% (N= 202; Figure 2). Of those owners who had used pheromone products (Adaptil® diffuser, spray or collar, N=316), 28.8% considered these to be effective. The reported success rates were 27.0% for nutraceuticals (N=211), 35.1% for herbal products (N=282), 31.2% for homeopathic products (N=250), 33.5% for Bach flowers (N=33.5) and 31.1% for essential oils (N=183), respectively.

**Figure 1.**
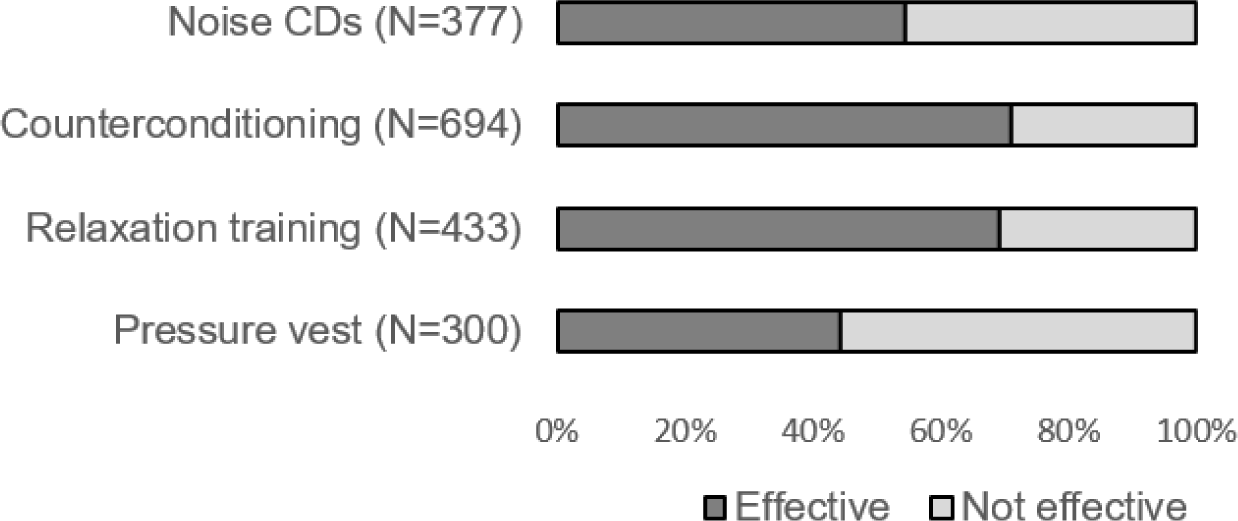
Owner-reported success rates of different behavior modification approaches and pressure vests.

**Figure 2.**
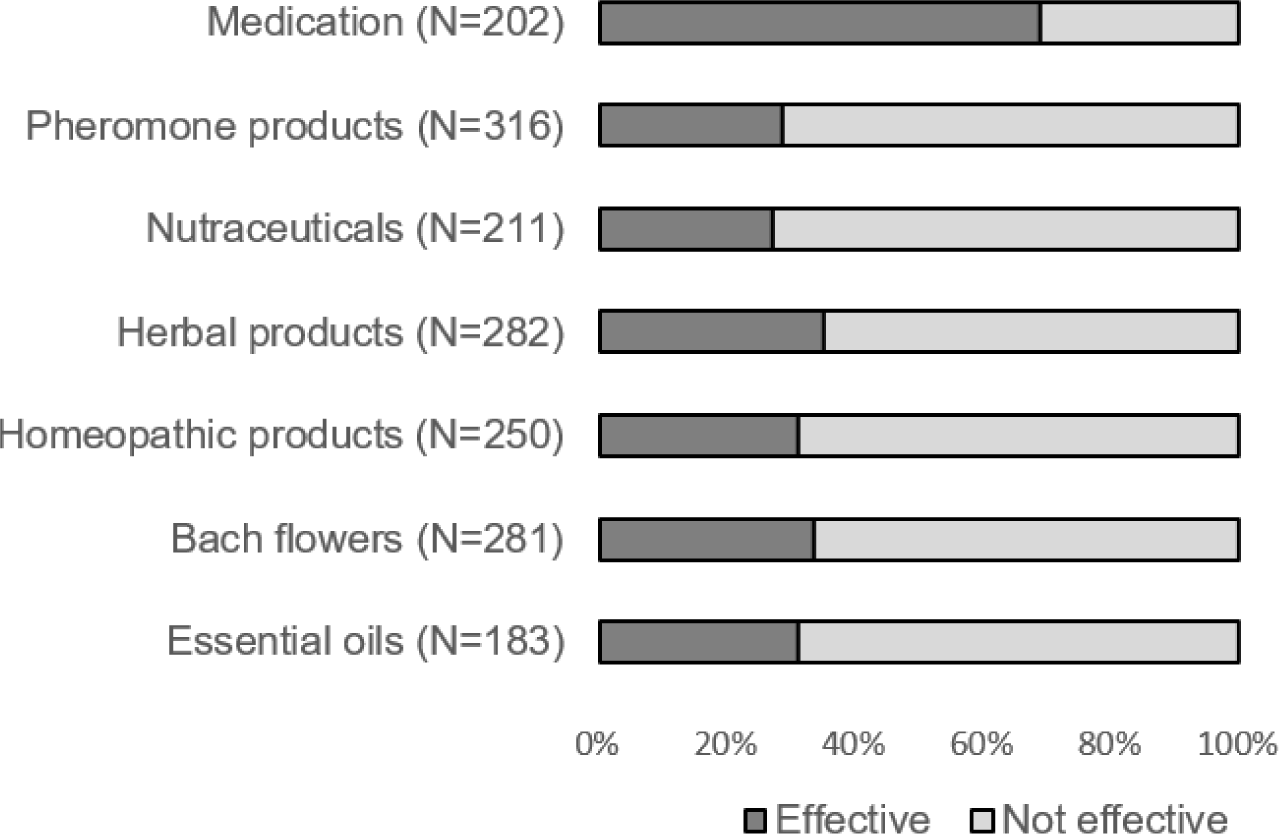
Owner-reported success rates of prescription medication, pheromones, calming nutraceuticals, herbal remedies, homeopathic products, Bach flowers and essential oils.

Since the question on which specific products were used were optional, data on the type of prescription medication used were available for 103 dogs. Of those, 86 received one type of drug, 13 received a combination of two drugs and four received three drugs. Alprazolam (N=32) and Dexmedetomidine (N=19) were the most frequently prescribed single drugs. The most frequent combination was Trazodone & Alprazolam (N=7). Other combinations included Dexmedetomidine combined with Diazepam or Alprazolam. Table 4 shows the reported effectiveness of the drugs in the sample, bearing in mind that the sample size was very small for most types of medication.

**Table 4.** Owner-reported effectiveness of different prescription medications

|  | N | Effective | Effective (%) |
| --- | --- | --- | --- |
| Alprazolam | 32 | 29 | 90.63 |
| Dexmedetomidine | 19 | 14 | 73.68 |
| Diazepam | 8 | 4 | 50 |
| Trazodone | 8 | 6 | 75 |
| Acepromazine | 4 | 4 | 100 |
| Benadryl | 3 | 3 | 100 |
| Fluoxetine | 3 | 3 | 100 |
| Clomipramine | 2 | 2 | 100 |
| Clonidine | 2 | 0 | 0 |
| Sertraline | 1 | 1 | 100 |
| Tramadol | 1 | 1 | 100 |
| >1 drug | 17 | 13 | 76.47 |
| Trazodone & Alprazolam | 7 | 7 | 100 |

## Discussion

An online survey was performed to explore how owners managed potential firework fears in their dogs and which interventions they perceived as effective in improving dogs’ firework fears.

### Management

In order to gauge whether management measures can affect the progression of firework fears in dogs, the four management factors were correlated with the Fear progression score. The results indicated that Environmental modification” (providing a hiding place, keeping windows and blinds shut, and playing music) was associated with a significant deterioration of firework fears, whereas provision of chews, play and food during fireworks in general, as well as deliberately providing these rewards when loud bangs occurred (principal component “Feed/Play”) was associated with a significant improvement. No association was found between fear progression and the “Alternative” component (use of nutraceuticals, pheromones, herbal products, homeopathic products, Bach flowers, and essential oils) or the “Interaction” component (allowing body contact, petting and talking to the dog when loud bangs occurred).

The positive correlation between Fear progression and “Environmental Modification” is unlikely to indicate that measures of environmental modification were *contributing* to increased fear severity. Conversely it is probable that owners who perceived an increased fear response in their dogs were more likely to take measures of environmental modification, since these are commonly recommended measures for dogs fearful of fireworks (e.g. Mills et al., 2003; Pike et al., 2015; Sherman and Mills, 2008). Of real interest, however, is the significant negative association of Fear progression and the “Feed/Play” component. It is reasonable to assume that this reflects a positive effect of the intervention on fear progression rather than vice versa, since owners experiencing an improvement in their dogs’ fears should be less likely and not more likely to attempt a given intervention. It appears that using appetitive stimuli during firework exposure not only helped to keep the fear at bay during the event, but may even have contributed to a longer-lasting improvement in fearfulness, possibly exerting some counterconditioning effects even without a systematic desensitization/ counterconditioning approach. Alternatively, it is possible that dog owners who used food and toys during fireworks also performed ad-hoc counterconditioning (providing these incentives contingent on occurrence of noises) in everyday life.

A common recommendation is to ignore the dog when it shows fearful behavior (e.g. Mills et al., 2003; Pike et al., 2015; Sherman and Mills, 2008). The current study indicated that interacting with the dog is neither associated with improvement nor with deterioration of the dogs’ fear level over time. Similarly, Dreschel and Granger (2005) found no influence of the owner’s behavior on behavior and cortisol levels in dogs after a playback of a thunderstorm recordings. Whether interacting with a fearful dog has beneficial or detrimental effects on fear level in the firework situation itself requires further investigation. There is, however, evidence from other stressful situations that being petted and talked to is associated with lower physiological or behavioral stress indicators in dogs (e.g. Csoltova et al., 2017; Hennessy et al., 1998; Lynch and McCarthy, 1967), and some physiological mechanisms for this have been identified (e.g. Handlin et al., 2011; Kostarczyk and Fonberg, 1982; Odendaal and Meintjes, 2003; Rehn et al., 2014). Thus, there might be potential beneficial effects of attending to fearful dogs also in firework situations, even if it cannot be ruled out that this reinforces some behavioral actions, such as attention-seeking, by instilling some fear relief (negative reinforcement).

### Behavior modification and training

The above-mentioned correlation between higher scores on the “Feed/Play” principal component and a more favorable fear progression has indicated that using food and play during fireworks is an effective way of alleviating noise fears in dogs. This is also confirmed by the owners’ own perceptions: Counterconditioning (defined in the questionnaire as “for example, giving a treat or playing with the dog after a noise has occurred”) was considered to be effective by 70.8% of those who had tried this technique. It is commonly believed that desensitization/ counterconditioning can only be successful in achieving a lasting emotional/ behavioral change if pets are kept under their threshold or tolerance level (Horwitz and Pike, 2014, but see a critical evaluation of this assumption in Klein, 1969; Wilson and Davison, 1971). While this would clearly be the ideal approach, the results of the current study indicate that even ad-hoc, non-systematic, counterconditioning in everyday life (such as giving a treat, playing with the dog or celebrating a little ‘party’) whenever a loud noise occurs, as well as during firework events, can contribute to an improvement of firework fears in dogs. While some dogs may be too stressed to eat during exposure to loud noises, providing high-value food after any noise occurrence is a simple strategy that can be easily implemented also by inexperienced owners and was actually used by over half the respondents in this study (N=694). Using this technique as a preventative measure appears to be extremely effective in preventing the development of firework fears in the first place (Riemer, 2019). This method is thus highly recommendable in both the prevention and treatment of noise fears in dogs.

Interestingly, with a success rate of a 69.3% according to the owners, relaxation training was similarly effective as counterconditioning in alleviating firework fears in dogs. Although relaxation was an inherent component of the original concept of “systematic desensitization” in people sensu Wolpe, (1958, reviewed in Thomas et al., 2012; Wilson and Davison, 1971), relaxation training is less prominently mentioned in the current literature on clinical animal behavior. Nonetheless, a relatively high number of respondents in the sample (N=433) had attempted this technique. The current questionnaire did not go into detail about the type of training protocols that were used, i.e. whether a classically conditioned approach was mostly used or whether respondents achieved a relaxed state through positive reinforcement of relaxed behaviors. Further research is therefore warranted on the optimal relaxation protocols. Similarly as with counterconditioning, it appears that respondents did not necessarily use relaxation training in a systematic way with a gradual increase in stimulus intensity; nonetheless, the technique seems to have been successful in a large proportion of cases, at least according to the owners’ perceptions.

The current gold standard and most studied behavioral approach in the therapy of noise fears is to perform desensitization/ counterconditioning (DSCC) using noise recordings (e.g. Horwitz and Mills, 2012; Levine et al., 2007; Levine and Mills, 2008; Mills et al., 2003; Sherman and Mills, 2008). Nonetheless, at 54.4% reported effectiveness, the use of noise recordings had a considerably lower success rate than ad-hoc counterconditioning and relaxation training in the current study. The rate of success is also somewhat lower than reported in previous studies on the use of noise CDs, possibly reflecting the more long-term view. For example, in a follow-up study one year after the first implementation of DSCC using noise CDs, 66% of owners indicated either “a moderate or great improvement” in their dog’s fear of fireworks, even though only few of the respondents had continued to use the CD recording (Levine and Mills, 2008), and reported success rates were even higher in the short-term (Levine et al., 2007; Sherman and Mills, 2008).

Perhaps owners who received the CD from professionals in veterinary behavior medicine were more motivated to follow through the procedure, or they had received better instructions on how to approach DSCC – although Mills et al. (2003) and Levine et al. (2007) found that many owners did not necessarily adhere to the instructions very well, and that two CDs with different types of instructions appeared to be similarly effective. The published studies had relatively high dropout rates. While personal reasons were mostly given (Levine et al., 2007), it cannot be ruled out that those owners who perceived no success of treatment were more likely to discontinue and were thus not available for reporting on the perceived success (or lack thereof) at the study end. Also with relatively small sample sizes in the cited studies compared to the current one, more random variation can be expected in the former. Additionally, treatment success might be over-reported when personal follow-up via phone calls are made as in Levine et al. (2007), Levine and Mills (2008) and Mills et al. (2003), since it is suggested that patients may report positive outcomes to the treating clinician out of politeness, enhancing the placebo effect (Kienle and Kiene, 1997). It is furthermore possible that the advice to introduce a “safe haven” (“a location in the home in which the dog had only positive experiences – not the same location to which the dog normally hid when fearful” (Levine et al., 2007) and the provision of Dog Appeasing Pheromone (Levine et al., 2007) enhanced the treatment success compared to the use of CD recordings alone (c.f. Mills et al., 2003).

Some difficulties with using noise recordings for DSCC have been recognized. They may not be realistic enough, depending on the quality of the recording or the speakers, the setup of the room influencing acoustics, and lacking associated stimuli such as flashes (Sheppard and Mills, 2003; Shull-Selcer and Stagg, 1991). Even under optimal conditions, some dogs will not respond to the simulations (Shull-Selcer and Stagg, 1991). Moreover, DSCC using noise recording is a time-consuming process, and may need to be repeated periodically to prevent a relapse, which some owners may be unwilling or unable to provide (Levine and Mills, 2008; Sheppard and Mills, 2003).

Thus, while noise recordings clearly have a place in the treatment of noise fears, ad-hoc counterconditioning to noises and relaxation training should complement this treatment strategy. In particular, using treats or other incentives whenever a (loud) noise occurs may be an easier strategy to implement in everyday life for many owners, as unlike both DSCC with noise recordings and relaxation training, it requires no targeted training other than having some food on hand – although clearly, the more structured training can be done, the better the outcome is likely to be.

### Products

While the reported success rates for counter-conditioning and relaxation training were relatively high, behavioral methods on their own may often not be sufficient to provide adequate fear relief. Accordingly, a wide range of products with putatively calming properties are on the market; however, for most of these, there is a lack of good-quality evidence on their effectiveness.

In the current study, the highest success rates of all product categories was achieved for prescription medication, even though it can be presumed that dogs that were medicated were affected by more severe noise fears than those receiving no medication. Since medications undergo a rigorous testing process (albeit not necessarily for the indication “noise fears”) before they are licensed, and work at the level of the central nervous system, this should be expected. While the overall sample size for users of medication was large in the current study (N=202), only a smaller number of owners had indicated which medications they had used. Of primary interest here are the two most commonly reported medications, Alprazolam (N=32) and Dexmedetomidine (N=19). The success rate was very high for Alprazolam at over 90%, while Dexmedetomidine was considered to be effective in over 73% of cases. The latter figure is very similar to the published effectiveness of Dexmedetomidine in a placebo-controlled study, in which a good or excellent treatment effect was found in 72% in the dexmedetomidine group, compared to 37% in the placebo group (Korpivaara et al., 2017). Too few respondents have named other types of medications so that no definitive conclusions can be drawn regarding their effectiveness.

Overall, pharmacological treatment is no doubt recommendable for dogs with severe noise fears, both as a management strategy to help them cope in the case of firework events and possibly to complement training efforts and facilitate learning during behavior modification treatment. Many owners, however, view medication only as a last resort and prefer to use more “natural” treatments, due to concerns such as potential side effects (Notari and Gallicchio, 2008; Sheppard and Mills, 2003). Unfortunately, very few of these products have demonstrated any evidence of effectiveness (Landsberg et al., 2015b).

Remarkably, all types of ‘alternative’ products in the current survey – including herbal products, nutraceuticals, homeopathic remedies, essential oils, but also pheromones – had reported success rates in the range of 27 to 35 percent. This is approximately the rate of success that would be expected based on a placebo effect. A placebo is “an inert medication used for its psychological effect, or for purposes of comparison in an experiment”, with a placebo effect defined as “any improvement or change in subjective discomfort or illness resulting from an intervention possessing no physical effect” (Tavel, 2014). When other individuals (clinicians or caregivers) rate the effectiveness of a given intervention, we may observe a caregiver placebo effect, which often has even larger effect sizes than those reported by the patients themselves (Rief et al., 2009). A caregiver effect in companion animals has also been defined as “improved ratings of outcomes in companion animals in the absence of improvement in objective measures” (Gruen et al., 2017). Whether a placebo-by-proxy effect exists in veterinary medicine – meaning that the caregiver’s belief alters their interaction with the pet which consequentially has a true beneficial effect on the animal – is not known (Gruen et al., 2017). Instead it is concerning that many interventions leading to better welfare according to the owners’ subjective assessments may in fact have no benefit on the animal at all.

In the medical literature, response rates to placebos are commonly in the range of around 35% (Muñana et al., 2010). In companion animals, caregiver placebo effects of up to 50-70% have been reported (Gruen et al., 2017). In the study by (Korpivaara et al., 2017) on the effect of dexmedetomidine vs placebo on firework fears in dogs, 37% of dogs in the placebo groups had a good or excellent treatment effect as judged by their owners. In the study by Cracknell and Mills (2008) evaluating the effect of a homeopathic remedy on firework fears in dogs, the reported rates of improvement did not differ significantly between treatment groups but were extremely high in both the placebo group (65%) and the treatment group (71%), although the fact that simple behavioral advice had also been given might have contributed to these high success rates in both groups (Cracknell and Mills, 2008). In a study on fluoxetine in the treatment of canine separation anxiety, global separation anxiety scores in the placebo-treated dogs improved in 44-51.3% of cases across the six weeks of treatment (Landsberg et al., 2008).

Thus, it seems likely that a placebo effect accounts for the perceived effectiveness of those products where the success rates were 35% or less in the current study, which was the case for pheromone products, nutraceuticals, herbal remedies, essential oils, as well as homeopathic remedies. In this context it is also notable that the many promising open-label trials have not been followed up by placebo controlled ones. Possibly this reflects a publication bias, with positive results from open-label trials potentially being more likely to be published than negative results in controlled studies.

Of the above-mentioned products, only Dog Appeasing Pheromone (Adaptil®) has been investigated in several studies on fear and anxiety in dogs. Some open-label trials reported high treatment success rates, but these were often confounded by the parallel application of the pheromones and behavioral management or CCSD training (Levine et al., 2007; Levine and Mills, 2008; Sheppard and Mills, 2003). One study suggested that using a combination of CDs and DAP resulted in better outcomes than when only one of these treatments was used (Mills et al., 2003). However, the high success rates for placebo treatments observed in placebo-controlled studies on treatments for firework fears in dogs (Cracknell and Mills, 2008; Korpivaara et al., 2017) call for a cautious interpretation of open-label studies. One parallel-group placebo controlled study using objective behavioral measures did indicate some beneficial effects of DAP collars on global and active fear scores during exposure to thunderstorm recordings in laboratory beagles. On the other hand, DAP treated dogs also spent more time hiding, so the results are not entirely conclusive, and the sample size was small (12 beagles per treatment group, Landsberg et al., 2015a). A systematic review on pheromone application in dogs and cats concluded that only a single study yielded sufficient evidence for a reduction in fear and anxiety by DAP in dogs, while six studies yielded insufficient evidence (Frank et al., 2010). Thus, also in view of the present study’s results, there is currently little evidence to support the effectiveness of dog appeasing pheromones as a calming agent in dogs fearful of fireworks. Similarly, no recommendations can be made for the use of nutraceuticals, herbal remedies, essential oils, and homeopathic products. Of course, perhaps with the exception of pheromones where the same ingredient, Dog Appeasing Pheromone, formed the basis of all products (diffusers, sprays or collars), the study does not allow drawing conclusions for a specific product in this case, as all products in a category were analyzed together.

Finally, the reported effectiveness for pressure vests (thundershirt, anxiety wrap) was 44%. Although this could still be in a range where a placebo effect is possible, the proportion of respondents considering pressure vests effective was considerably higher compared to all the ‘alternative’ products, indicating that pressure vests may potentially have beneficial effects in some dogs. Further research is needed, as previous studies indicated a possible small beneficial effect but were not conclusive either (reviewed in Buckley, 2018).

### Critical evaluation

Like most studies assessing the effectiveness of treatments for noise fears in dogs (e.g. Cracknell and Mills, 2008; Crowell-Davis et al., 2003; Korpivaara et al., 2017; Levine et al., 2007; Levine and Mills, 2008; Sheppard and Mills, 2003), the current study relied on owners’ assessments of their dogs’ behavior. Therefore it cannot be avoided that the results are affected by the owners’ interpretation and may be subject to bias. However, studies indicated that owners’ ratings of their dogs’ fear are reliable and demonstrate external validity (e.g. Tiira and Lohi, 2014), although they are susceptible to caregiver placebo effects (e.g. Cracknell and Mills, 2008). Based on previous placebo-controlled studies, we know approximately in what range placebo effects can be expected, as discussed above. While randomized double-blind placebo-controlled studies, and using video analysis to assess dogs’ behavior, are no doubt the gold standard for assessing the effect of a given intervention, they are often limited by the costs and manpower required for both conducting and administrating the study. Additionally, ethical concerns might be raised (c.f. Frank et al., 2010). Using a large-scale questionnaire survey thus represents a useful approach to gather data from a large sample of participants. Moreover, a broad survey like the current one may be less likely than localized trials of specific products to attract volunteer participants with high expectations that might lead to an overestimate of effectiveness (c.f. Muñana et al., 2010), or to induce participants to over-report effectiveness of treatments out of politeness to the treating doctor (Kienle and Kiene, 1997). It can thus be viewed as an advantage that the prescribing clinicians and the researchers were not the same persons in the current study. Owners were furthermore able to draw from their long-term experiences (i.e. not only for a few weeks within a particular study). While recall biases cannot be ruled out, the possibility to test a given product over a longer time period or during several firework events may lead to a more accurate assessment than when owners are asked to make an assessment over only a few weeks.

By asking owners to simply rate a given intervention as “effective” or “not effective”, the outcome is relatively crude. Nonetheless it is useful, since effectiveness of products is often measured by the proportion in the population in which the treatment had an effect, and together with the large sample size, it allows us to compare relative effectiveness of different treatments – even if it does not allow conclusions in absolute numbers regarding the true effectiveness of the interventions.

## Summary

Owner-reported effectiveness of different treatment options for firework fears in dogs was analyzed from a large-scale questionnaire survey, with over 180 respondents per treatment category. Ad-hoc counter-conditioning (providing desirable stimuli after the occurrence of noises) was associated with a significant improvement in the severity of firework fears and was considered as effective by over 70% of owners. Relaxation training was reported to be almost as successful, while noise CDs were considered effective by only 55%. Forty-four percent of respondents deemed pressure vests to be effective. With prescription medication, improvements were noted by 70% of owners. While individual products were not evaluated, the reported success rates for the categories “pheromones”, “herbal products”, “nutraceuticals”, “essential oils”, “homeopathic remedies” and “Bach flowers” (all in the range of 27-35%) are not higher than would be expected based on a placebo effect. Thus, besides use of anxiolytic medication, counterconditioning and relaxation training appear to be the most effective strategies in the treatment of firework fears in dogs and should complement the standard behavioral technique of desensitization/ counterconditioning with noise recordings.

## Acknowledgements

Many thanks to all the dog owners who took the time to fill in this questionnaire and to everybody who helped to spread the survey. Further thanks go to Hanno Würbel and Alja Mazzini for feedback on the manuscript. S.R. was supported by an SNF Ambizione Grant Project PZ00P3_174221.

## Authorship statement

The idea for the paper was conceived by Stefanie Riemer. The questionnaire design, distribution, analysis and writing was performed by Stefanie Riemer.

## Data Availability

The raw data set can be accessed under https://figshare.com/articles/Riemer_2019_dogs_fireworks_data_xls/9657395.

## References

Blackwell, E.J., Bradshaw, J.W.S., Casey, R.A., 2013. Fear responses to noises in domestic dogs: Prevalence, risk factors and co-occurrence with other fear related behaviour. Appl. Anim. Behav. Sci. 145, 15–25.

Buckley, L.A., 2018. Are Pressure Vests Beneficial at Reducing Stress in Anxious and Fearful Dogs? Vet. Evid. 3.

Cottam, N., Dodman, N.H., Ha, J.C., 2013. The effectiveness of the Anxiety Wrap in the treatment of canine thunderstorm phobia: An open-label trial. J. Vet. Behav. Clin. Appl. Res. 8, 154–161.

Cracknell, N.R., Mills, D.S., 2008. A double-blind placebo-controlled study into the efficacy of a homeopathic remedy for fear of firework noises in the dog (*Canis familiaris*). Vet. J. 177, 80–88.

Crowell-Davis, S.L., Seibert, L.M., Sung, W., Parthasarathy, V., Curtis, T.M., 2003. Use of clomipramine, alprazolam, and behavior modification for treatment of storm phobia in dogs. J. Am. Vet. Med. Assoc. 222, 744–748.

Csoltova, E., Martineau, M., Boissy, A., Gilbert, C., 2017. Behavioral and physiological reactions in dogs to a veterinary examination: Owner-dog interactions improve canine well-being. Physiol. Behav. 177, 270–281.

DePorter, T.L., Landsberg, G.M., Araujo, J.A., Ethier, J.L., Bledsoe, D.L., 2012. Harmonease Chewable Tablets reduces noise-induced fear and anxiety in a laboratory canine thunderstorm simulation: a blinded and placebo-controlled study. J. Vet. Behav. Clin. Appl. Res. 7, 225–232.

Dreschel, N.A., Granger, D.A., 2005. Physiological and behavioral reactivity to stress in thunderstorm-phobic dogs and their caregivers. Appl. Anim. Behav. Sci. 95, 153–168.

Engel, O., Müller, H.W., Klee, R., Francke, B., Mills, D.S., 2019. Effectiveness of imepitoin for the control of anxiety and fear associated with noise phobia in dogs. J. Vet. Intern. Med. 33, 2675–2684.

European_Medicines_Agency, 2015. Sileo Dexmedetomidine hydrochloride – EPAR summary for the public.

Fish, R.E., Foster, M.L., Gruen, M.E., Sherman, B.L., Dorman, D.C., 2017. Effect of Wearing a Telemetry Jacket on Behavioral and Physiologic Parameters of Dogs in the Open-Field Test. J. Am. Assoc. Lab. Anim. Sci. 56, 382–389.

Frank, D., Beauchamp, G., Palestrini, C., 2010. Systematic review of the use of pheromones for treatment of undesirable behavior in cats and dogs. J. Am. Vet. Med. Assoc. 236, 1308–1316.

Gruen, M.E., Case, B.C., Foster, M.L., Lazarowski, L., Fish, R.E., Landsberg, G., DePuy, V., Dorman, D.C., Sherman, B.L., 2015. The use of an open-field model to assess sound-induced fear and anxiety-associated behaviors in Labrador retrievers. J. Vet. Behav. 10, 338–345.

Gruen, M.E., Dorman, D.C., Lascelles, B.D.X., 2017. Caregiver Placebo Effect in Analgesic Clinical Trials for Painful Cats with Naturally-Occurring Degenerative Joint Disease. Vet. Rec. 180, 473.

Gruen, M.E., Sherman, B.L., 2008. Use of trazodone as an adjunctive agent in the treatment of canine anxiety disorders: 56 cases (1995--2007). J. Am. Vet. Med. Assoc. 233, 1902–1907.

Handlin, L., Hydbring-sandberg, E., Nilsson, A., Ejdebäck, M., Jansson, A., Uvnäs-moberg, K., 2011. Short-Term Interaction between Dogs and Their Owners□: Effects on Oxytocin, Cortisol, Insulin and Heart Rate — An Exploratory Study. Anthrozöos 24, 301–315.

Hennessy, M.B., T Williams, M., Miller, D.D., Douglas, C.W., Voith, V.L., 1998. Influence of male and female petters on plasma cortisol and behaviour: can human interaction reduce the stress of dogs in a public animal shelter? Appl. Anim. Behav. Sci. 61, 63–77.

Herron, M.E., Shofer, F.S., Reisner, I.R., 2008. Retrospective evaluation of the effects of diazepam in dogs with anxiety-related behavior problems. J. Am. Vet. Med. Assoc. 233, 1420–1424.

Horwitz, D., Mills, D., 2012. BSAVA manual of canine and feline behavioural medicine, second. ed. BSAVA.

Horwitz, D.F., Pike, A.L., 2014. Common sense behavior modification: a guide for practitioners. Vet. Clin. Small Anim. Pract. 44, 401–426.

Kienle, G.S., Kiene, H., 1997. The powerful placebo effect: fact or fiction? J. Clin. Epidemiol. 50, 1311–1318.

Klein, B., 1969. Counterconditioning and fear reduction in the rat. Psychon. Sci. 17, 150–151.

Korpivaara, M., Laapas, K., Huhtinen, M., Schöning, B., Overall, K., 2017. Dexmedetomidine oromucosal gel for noise-associated acute anxiety and fear in dogs—a randomised, double-blind, placebo-controlled clinical study. Vet. Rec. 180, 356.

Kostarczyk, E., Fonberg, E., 1982. Heart rate mechanisms in instrumental conditioning reinforced by petting in dogs. Physiol. Behav. 28, 27–30.

Landsberg, G., Beck, A., Lopez, A., Deniaud, M., Araujo, J., Milgram, N., 2015a. Dog-appeasing pheromone collars reduce sound-induced fear and anxiety in beagle dogs: a placebo-controlled study. Vet. Rec. 177, 260.

Landsberg, G., Mougeot, I., Kelly, S., Milgram, N., 2015b. Assessment of noise-induced fear and anxiety in dogs: modification by a novel fish hydrolysate supplemented diet. J. Vet. Behav. 10, 391–398.

Landsberg, G.M., Melese, P., Sherman, B.L., Neilson, J.C., Zimmerman, A., Clarke, T.P., 2008. Effectiveness of fluoxetine chewable tablets in the treatment of canine separation anxiety. J. Vet. Behav. 3, 12–19.

Levine, E.D., Mills, D.S., 2008. Long-term follow-up of the efficacy of a behavioural treatment programme for dogs with firework fears. Vet. Rec. 162, 657–9.

Levine, E.D., Ramos, D., Mills, D.S., 2007. A prospective study of two self-help CD based desensitization and counter-conditioning programmes with the use of Dog Appeasing Pheromone for the treatment of firework fears in dogs (*Canis familiaris*). Appl. Anim. Behav. Sci. 105, 311–329.

Linting, M., Meulman, J.J., Groenen, P.J.F., van der Koojj, A.J., 2007. Nonlinear principal components analysis: introduction and application. Psychol. Methods 12, 336.

Linting, M., van der Kooij, A., 2012. Nonlinear principal components analysis with CATPCA: a tutorial. J. Pers. Assess. 94, 12–25.

Lynch, J.J., McCarthy, J.F., 1967. The effect of petting on a classically conditioned emotional response. Behav. Res. Ther. 5, 55–62.

McMillan, F.D., 2019. Mental health and well-being in animals, 2nd ed. CABI.

Mills, D., 2005. Management of noise fears and phobias in pets. In Pract. 27, 248.

Mills, D., Dubé, M.B., Zulch, H., 2012. Principles of pheromonatherapy. Stress Pheromonatherapy Small Anim. Clin. Behav. 127–145.

Mills, D.S., 2003. Medical paradigms for the study of problem behaviour: a critical review. Appl. Anim. Behav. Sci. 81, 265–277.

Mills, D.S., Estelles, M.G., Coleshaw, P.H., Shorthouse, C., 2003. Retrospective analysis of the treatment of firework fears in dogs. Vet. Rec. 153, 561–562.

Muñana, K.R., Zhang, D., Patterson, E.E., 2010. Placebo effect in canine epilepsy trials. J. Vet. Intern. Med. 24, 166–170.

Newall, C., Watson, T., Grant, K.-A., Richardson, R., 2017. The relative effectiveness of extinction and counter-conditioning in diminishing children’s fear. Behav. Res. Ther. 95, 42–49.

Notari, L., Gallicchio, B., 2008. Owners’ perceptions of behavior problems and behavior therapists in Italy: A preliminary study. J. Vet. Behav. 3, 52–58.

Odendaal, J.S.J., Meintjes, R.A., 2003. Neurophysiological correlates of affiliative behaviour between humans and dogs. Vet. J. 165, 296–301.

Ogata, N., Dodman, N.H., 2011. The use of clonidine in the treatment of fear-based behavior problems in dogs: an open trial. J. Vet. Behav. Clin. Appl. Res. 6, 130–137.

Orlando, J.M., 2018. Behavioral nutraceuticals and diets. Vet. Clin. Small Anim. Pract. 48, 473–495.

Overall, K., 2013. Manual of clinical behavioral medicine for dogs and cats. Elsevier, St Louis, MO.

Pekkin, A.-M., Hänninen, L., Tiira, K., Koskela, A., Pöytäkangas, M., Lohi, H., Valros, A., 2016. The effect of a pressure vest on the behaviour, salivary cortisol and urine oxytocin of noise phobic dogs in a controlled test. Appl. Anim. Behav. Sci. 185, 86–94.

Pike, A.L., Horwitz, D.F., Lobprise, H., 2015. An open-label prospective study of the use of l-theanine (Anxitane) in storm-sensitive client-owned dogs. J. Vet. Behav. 10, 324–331.

Rehn, T., Handlin, L., Uvnäs-Moberg, K., Keeling, L.J., 2014. Dogs’ endocrine and behavioural responses at reunion are affected by how the human initiates contact. Physiol. Behav. 124, 45–53.

Rief, W., Nestoriuc, Y., Weiss, S., Welzel, E., Barsky, A.J., Hofmann, S.G., 2009. Meta-analysis of the placebo response in antidepressant trials. J. Affect. Disord. 118, 1–8.

Riemer, S., 2019. Not a one-way road – severity, progression and prevention of firework fears in dogs. PLoS One 14, e0218150. https://doi.org/10.1371/journal.pone.0218150

Sheppard, G., Mills, D.S., 2003. Evaluation of dog-appeasing pheromone as a potential treatment for dogs fearful of fireworks. Vet. Rec. J. Br. Vet. Assoc. 152, 432–436.

Sherman, B.L., Mills, D.S., 2008. Canine anxieties and phobias: an update on separation anxiety and noise aversions. Vet. Clin. North Am. Small Anim. Pract. 38, 1081–1106.

Shull-Selcer, E.A., Stagg, W., 1991. Advances in the understanding and treatment of noise phobias. Vet. Clin. North Am. Small Anim. Pract. 21, 353–367.

Storengen, L.M., Lingaas, F., 2015. Noise sensitivity in 17 dog breeds: Prevalence, breed risk and correlation with fear in other situations. Appl. Anim. Behav. Sci. 171, 152–160.

Talegón, M.I., Delgado, B.A., 2011. Anxiety disorders in dogs, in: Anxiety Disorders. InTech, pp. 261–280.

Tavel, M.E., 2014. The placebo effect: the good, the bad, and the ugly. Am. J. Med. 127, 484–488.

Thomas, B.L., Cutler, M., Novak, C., 2012. A modified counterconditioning procedure prevents the renewal of conditioned fear in rats. Learn. Motiv. 43, 24–34.

Tiira, K., Lohi, H., 2014. Reliability and validity of a questionnaire survey in canine anxiety research. Appl. Anim. Behav. Sci. 155, 82–92.

Tiira, K., Sulkama, S., Lohi, H., 2016. Prevalence, comorbidity, and behavioral variation in canine anxiety. J. Vet. Behav. Clin. Appl. Res. 16, 36–44.

Wilson, G.T., Davison, G.C., 1971. Processes of fear reduction in systematic desensitization: Animal studies. Psychol. Bull. 76, 1.

Wolpe, J., 1958. Psychotherapy by reciprocal inhibition. Stanford University Press., Palo Alto.

